# Structure of a AAA+ unfoldase in the process of unfolding substrate

**DOI:** 10.1101/105866

**Authors:** Zev. A Ripstein, Rui Huang, Rafal Augustyniak, Lewis E. Kay, John L. Rubinstein

## Abstract

AAA+ unfoldases are thought to unfold substrate through the central pore of their hexameric structures, but how this process occurs is not known. VAT, the *Thermoplasma acidophilum* homologue of eukaryotic CDC48/p97, works in conjunction with the proteasome to degrade misfolded or damaged proteins. We show that in the presence of ATP, VAT with its regulatory N-terminal domains removed unfolds other VAT complexes as substrate. We captured images of this transient process by electron cryomicroscopy (cryo-EM) to reveal the structure of the substrate-bound intermediate. Substrate binding breaks the six-fold symmetry of the complex, allowing five of the six VAT subunits to constrict into a tight helix that grips an ~80 Å stretch of unfolded protein. The structure suggests a processive hand-over-hand unfolding mechanism, where each VAT subunit releases the substrate in turn before re-engaging further along the target protein, thereby unfolding it.

---

The 85 kDa VAT protein, which is composed of an N-terminal regulatory domain (NTD) and two tandem nucleotide-binding AAA+ domains (NBD1 and NBD2), assembles into a 500 kDa homohexamer (Fig. 1 Supplement 1). Truncation of the first 182 residues of VAT removes the regulatory NTD, producing a construct (ΔN-VAT) that has both higher ATPase and unfoldase activity than the intact protein ^1–3^. In solutions of ΔN-VAT, the enzyme itself is degraded in an ATP- and proteasome-dependent manner (Fig. 1a-c). The 20S proteasome does not hydrolyze ATP during proteolysis ^4^ and therefore this observation demonstrates that ΔN-VAT complexes must unfold other copies of ΔN-VAT via a process that requires ATP hydrolysis prior to degradation by the proteasome.

**Figure 1.**
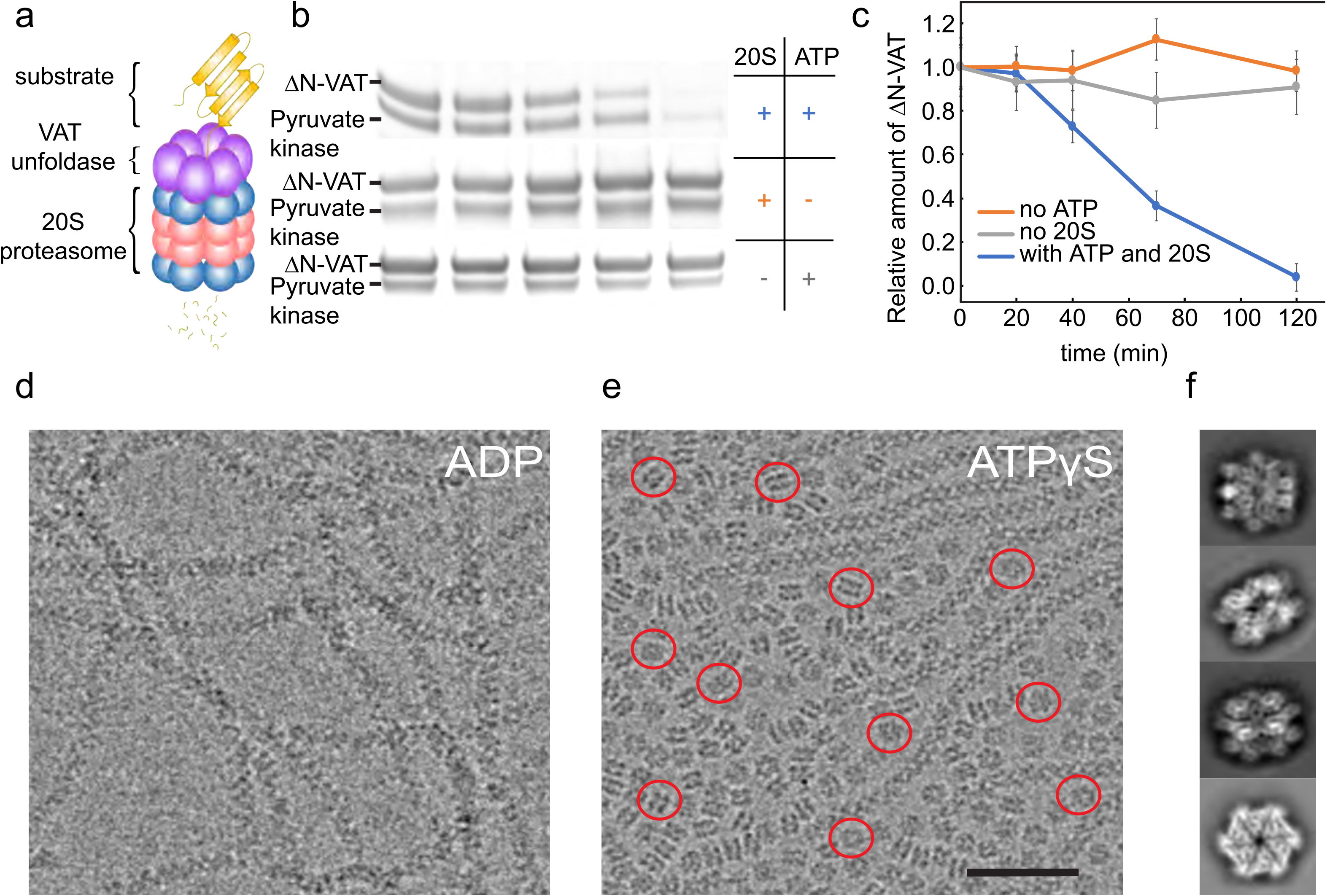
ΔN-VAT unfolds other copies of ΔN-VAT. **a,** Cartoon showing VAT (purple ring) working in concert with the 20S proteasome (blue and red rings) to degrade protein substrates (yellow) ^1^. **b,** An SDS-PAGE gel showing that both ΔN-VAT and pyruvate kinase (part of the ATP regenerating system) are degraded in an ATP and 20S proteasome dependent manner. **c,** Quantification of the self-unfolding reaction. All measurements done in triplicate. **d,** Representative micrographs of ΔN-VAT in the presence of 5 mM ADP. **e,** Representative micrographs of ΔN-VAT in the presence of 5 mM ATPγS. Example single particles are circled in red. Scale bar, 500 Å. **f,** 2D class average images showing views of single particles of ΔN-VAT in the presence of ATPγS.

Cryo-EM of full-length VAT previously showed that the enzyme adopts a helical split-ring conformation in the presence of ADP ^5^. ΔN-VAT particles with bound ADP assemble into long fibrils (Fig. 1d), presumably comprised of helical split-ring structures interacting with each other top-to-bottom. With the slowly-hydrolysable ATP-analogue ATPγS bound, full-length VAT exists in both the split-ring conformation and a six-fold symmetric stacked-ring conformation ^5^. Similarly, ΔN-VAT with bound ATPγS forms a mixture of fibrils (Fig. 1e) as well as single particles, the majority of which appeared to be six-fold symmetric (Fig. 1e, red circles). The fibrils could be excluded easily when selecting single particle images for analysis, allowing calculation of a cryo-EM map of the six-fold symmetric stacked-ring conformation of VAT at 3.9 Å resolution. This resolution enabled construction of an atomic model for the complex (Fig. 2, Fig. Supplement 1, Table 1). In the structure, a central pore runs along the six-fold symmetry axis of the complex through the center of the stacked NBD1 and NBD2 rings (Fig. 2a, left, hexagon). The stacked-ring structure of VAT resembles recent structures of the VAT homologue p97 in the fully ATP-loaded state ^6–8^, except for the orientations between the NBD1 and NBD2 domains in VAT monomers and the relative rotation of the NBD1 and NBD2 rings. All AAA+ unfoldases have conserved sequences that extend from the NBDs towards the center of the ring, forming ‘pore loops’ that are essential for unfolding substrate ^1^ (Fig. 2 Supplement 2). In the six-fold symmetric VAT structure, pore loop 1 of NBD1, containing conserved and essential tyrosine residues (Tyr264 and Tyr265) ^1^, forms an opening that is ~25 Å across while pore loop 2 of NBD1 (residues 299 to 305) forms a slightly tighter aperture that is ~18 Å across (Fig. 2 Supplement 2a, left). Similar apertures are produced by the corresponding pore loop 1 of NBD2 (containing conserved residues Trp 541 and Val 542) and pore loop 2 of NBD 2 (residues 575-579) (Fig. 2 Supplement 2a and b, left). An atomic model could be built for the entire map (Fig. 2b), including the linker region between NBD1 and NBD2 (Fig. 2c, left), and both NBD1 and NBD2 have strong density for ATPγS in their nucleotide-binding pockets (Fig. 2c, right) indicating that VAT adopts a rigid and symmetric conformation when fully loaded with nucleotide.

**Table 1.**
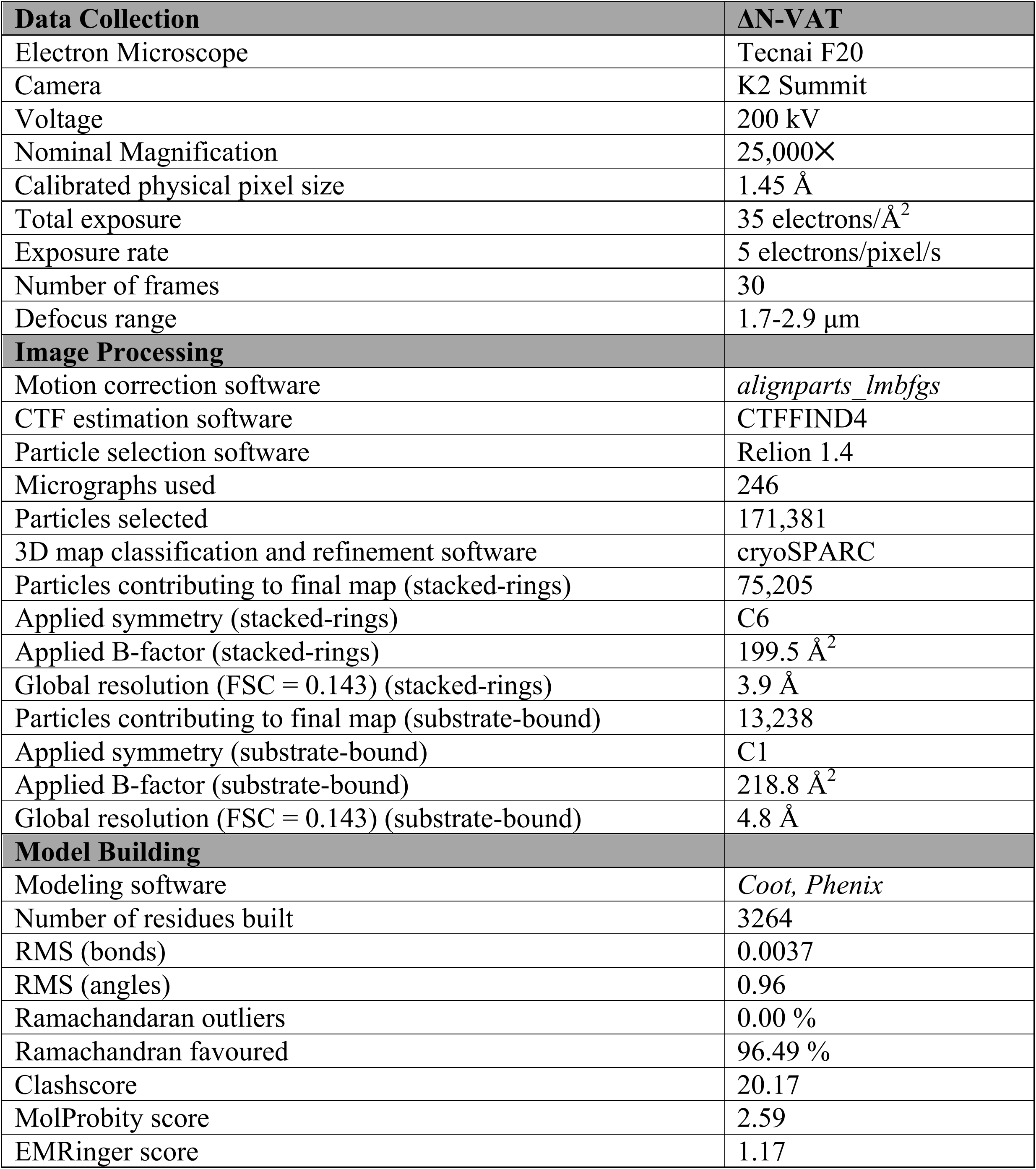
Cryo-EM data acquisition, processing, and atomic model statistics.

**Figure 2.**
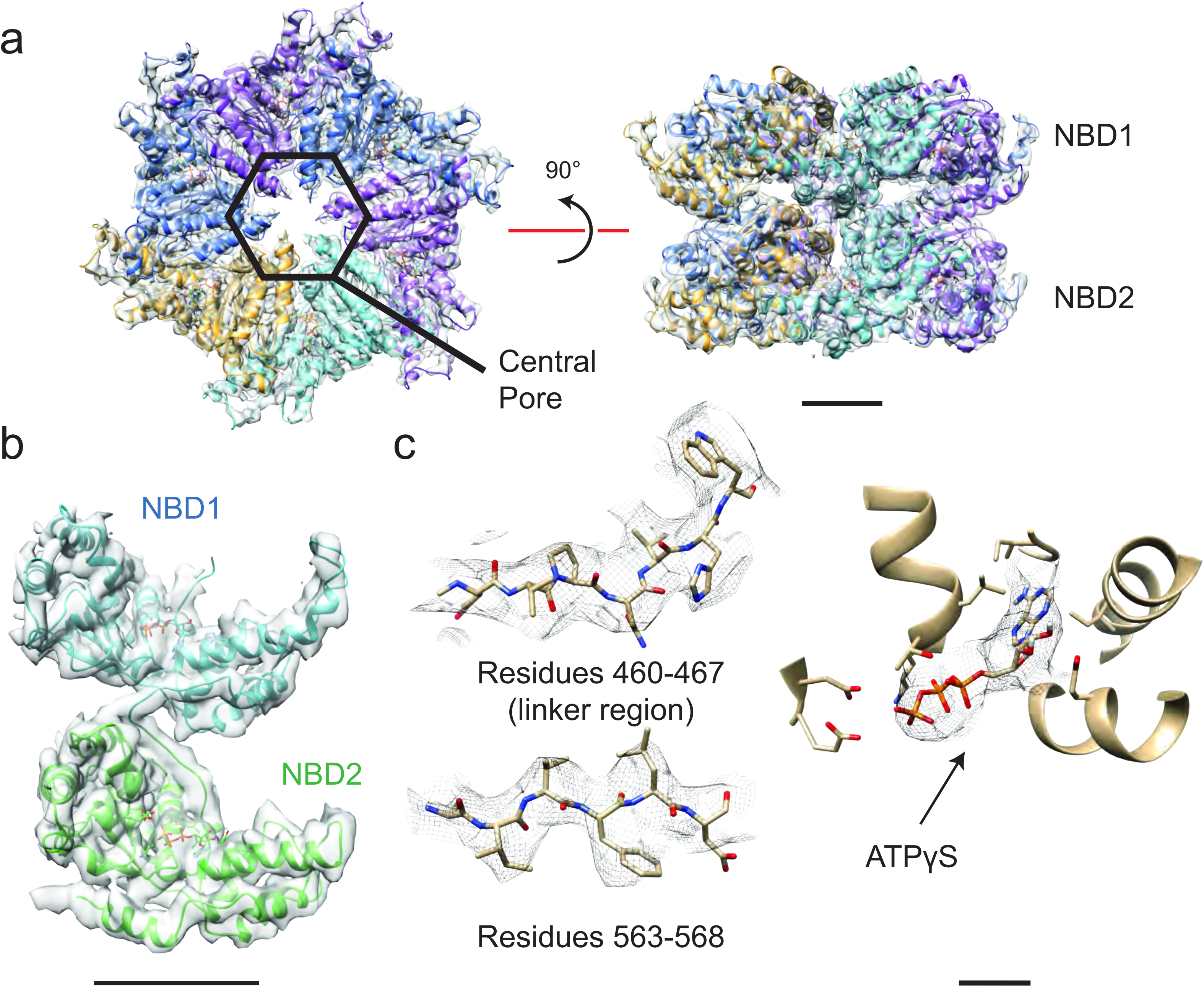
Structure of six-fold symmetric ΔN-VAT with bound ATPγS. **a,** Density map and atomic model for ATPγS-bound ΔN-VAT at 3.9 Å. A central pore runs through the middle of the complex. Scale bar, 25 Å. **b,** Density map and model for a single protomer of VAT shows both AAA+ domains and the linker between them. Scale bar, 25 Å. **c**, The cryo-EM density allowed identification of large amino acid side chains throughout ΔN-VAT, including in the linker region. Density corresponding to ATPγS is visible in all twelve catalytic sites. Scale bar, 5 Å.

Unsupervised classification of ATPγS-loaded ΔN-VAT particle images using a new *ab initio* 3D classification algorithm ^9^ allowed identification of a second conformation of the protein that was significantly different from known structures. The class corresponding to this conformation, containing ~15 % of particle images, showed two ΔN-VAT complexes in close apposition to one another (Fig. 3, Fig. 3 Supplement 1). Refinement of this structure to ~4.8 Å resolution revealed it to consist of a ΔN-VAT complex in the process of unfolding a neighboring complex as substrate (Fig. 3a). The map shows clear density for the substrate protein in an extended conformation as it runs almost directly through the central pore of the active ΔN-VAT (Fig. 3a and b, red density). Density for the substrate above NBD1 of the active ΔN-VAT complex is weak and fragmented, and is most clearly visualized without sharpening of the map (Fig. 3a, above dashed line) ^10^, suggesting that outside of the active enzyme the substrate has a flexible orientation relative to the pore of ΔN-VAT. The length of unfolded protein gripped by the pore of the active ΔN-VAT is ~80 Å, which corresponds to 12 to 14 extended amino acid residues.

In both NBD1 and NBD2 five of the six VAT subunits form close contacts with the substrate through their pore loops (Fig. 3b and c, and Fig. 2 Supplement 2a and b, middle). The hydrophobic residues of these pore loops create a tight helical sheath around the substrate (Fig. 3a, green). Breaking the six-fold symmetry of the complex forms a ‘seam’ where one protomer has interfaces that differ dramatically from other protomer-protomer contacts (Fig. 3c, gold) and the seam subunit does not interact with the substrate (Fig. 2 Supplement 2a and b, middle). This helical arrangement of the pore loop residues maximizes the contact area between the unfolded substrate and the pore loops of the five subunits that form the helix (Fig. 3b, Video 1). Unfolded substrate that spans the gap between the pore loops of NBD1 and NBD2 has a low density in the cryo-EM map, likely because of flexibility of the substrate in this region. The transition from the stacked-ring to the substrate-engaged conformation of ΔN-VAT occurs with rearrangement of NBD1 relative to NBD2 in each monomer and a rotation of the NBD1 ring against the NBD2 ring (Fig. 3 Supplement 2, Video 1 and 2). The two rings are also displaced horizontally relative to each other in the substrate-bound state and the central pore is tilted so that it passes between the rings at an angle of ~15° (Fig. 3 Supplement 2c). This rotation and displacement brings the NBD1 and NBD2 rings closer together than in the stacked-ring conformation, decreasing the distance between their pore loops.

**Figure 3.**
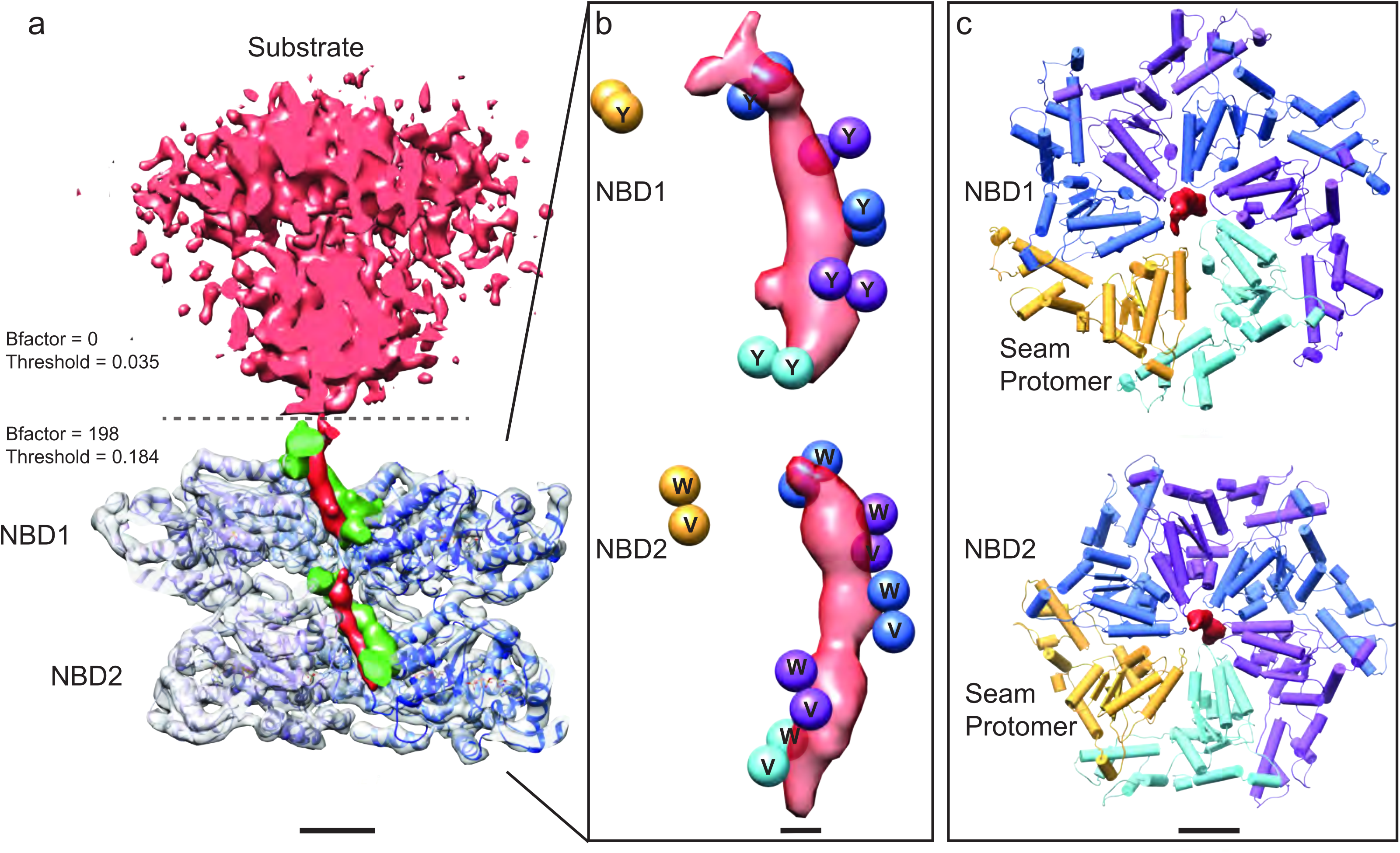
Substrate engagement by ΔN-VAT. **a,** Cut-away side view of the density map and model for ΔN-VAT with substrate bound. The substrate complex is shown unsharpened in pink above the dashed line. Density for the substrate-gripping loop residues is shown in green, with the enzyme-bound portion of the substrate shown as density (red) running through the center of the complex. Scale bar, 25 Å. **b,** Positions of the Catoms of the pore loop residues (spheres) surrounding the experimental density for substrate (red). Five of the six protomers contact the substrate. Scale bar, 5 Å. **c,** NBD1 and NBD2 rings with experimental density for substrate. Both NBD rings adopt similar conformations and a helical architecture, with five of the pore loops contacting the substrate. Scale bar, 25 Å.

In this study, only the newly identified conformation described above is found with substrate bound. Consequently, the six-fold symmetric stacked-ring state with ATPγS bound and the split-ring state with ADP bound ^5^ likely correspond to pre- and post-unfolding conformations of the enzyme. In the substrate-bound conformation, the seam protomer is positioned such that it does not interact with the substrate. Notably, recent structures of the 26S proteasome ^11,12^ also showed that either the Rpt5 or Rpt6 subunit in the hexameric Rpt ring of the 19S regulatory particle base is displaced from the pore, similar to the seam protomer in VAT (Fig. 3). Furthermore, the translocation mechanisms of AAA+ DNA helicases such as E1, Rho, and MCM, also involve a helical gripping state in which only five of six protomers interact with a single DNA strand ^13–15^. The structure presented here suggests an analogous substrate translocation mechanism for VAT where the enzyme operates processively with a ‘hand-over-hand’ mechanism, shown schematically in Figure 4a, focusing on protomer 1 outlined in green. In the first step of this cycle (Fig. 4a, top) protomer 1 is bound to unfolded substrate at the bottom of the helix of VAT protomers, protomer 2 in the seam position does not contact the substrate (Fig. 4a, gold), and protomer 3 binds substrate at the top of the helix near the remaining folded protein. The nucleotide binding sites shared by the seam protomer and protomer 1 are empty (see below). As the seam protomer reengages with substrate at the top of the helix (step 2: Fig. 4a, middle), protomer 1 disengages from the substrate and becomes the seam protomer, with protomer 6 now furthest along the unfolded substrate. The process repeats in a cyclical manner for additional protomers (e.g. step 3: Fig. 4a, bottom, Video 3). The power stroke for substrate unfolding is provided by ATP hydrolysis, which occurs in sequence around the VAT ring, with ADP released by the protomer adjacent to the seam (counter clockwise from the seam when viewed from NBD1 toward NBD2, Fig. 4b). Although the precise position where ATP hydrolysis and ADP release occur in the cycle is not apparent from our data, it is clear that ATP must rebind to the seam protomer as it engages with substrate at the top of the protomer helix. As the cycle continues, each protomer pulls a single section of the substrate, corresponding to ~13 Å or approximately two extended residues, through the central pore until it becomes the lowest protomer in the helix and releases the substrate (Video 3). The cyclic nature of the ATP hydrolysis gives directionality to the substrate translocation, allowing unfolded protein to be passed to the proteasome for degradation.

The cyclic ‘hand-over-hand’ model requires that each protomer hydrolyze ATP in turn (Fig. 4b). A single inactivated subunit would consequently have a severe effect on the rate of ATP hydrolysis, potentially stopping unfolding. To test the model, we measured the ATPase activity of VAT complexes that had been doped with different fractions (*r*) of catalytically-inactive protomers bearing mutations in the Walker B motifs of both NBD1 and NBD2 (E291Q, E568Q)^16^ relative to wild type VAT ATPase activity (*A*). These experiments were carried out in the presence of a GFP substrate with an 11-residue ssRA tag as substrate ^4^. For multimeric enzymes where each subunit functions independently, ATPase activity is expected to decrease linearly as increasing amounts of catalytically dead mutant are incorporated randomly into the oligomers (Fig. 4c, blue line). In contrast, the decrease in ATPase activity of the mixed complexes as a function of increasing concentration of catalytically dead mutant closely follows the *A* = (1 - *r*)^6^ curve expected for the proposed hand-over-hand model (Fig. 4c, gray line). The activity of VAT with increasing *r* decreases faster than the *A* = (1 - *r*)^6^ + 5*r*(1 - *r*)^5^ curve (Fig. 4c, dashed gray line) that is expected if ATP hydrolysis only stops when two of the six VAT monomers are inactivated.

The slight elevation relative to the *A* = (1 - *r*)^6^ profile in Figure 4c may reflect an ‘escape’ mechanism that allows stalled VAT complexes to disengage and then reengage with substrate, as seen with other molecular machines ^16^. In the absence of an escape mechanism ATP hydrolysis would proceed from one protomer to the next in a cyclical manner (Fig. 4b) until a defective subunit is reached, preventing further hydrolysis and, thus, subsequent substrate unfolding. In contrast, if the machine is able to disengage at this point it would be possible for other functional subunits to hydrolyze ATP, although at reduced rates. The existence of an escape mechanism therefore has important implications for substrate unfolding, even in the context of wild type VAT, by allowing the release of stalled substrate. This release minimizes both the loss of active enzyme and increases catalytic efficiency. Our ATPase assay data in Fig. 4c. cannot distinguish between a model where each protomer hydrolyzes ATP simultaneously or a sequential binding/hydrolysis mechanism as described in Fig. 4a and b. However, previous ATPase measurements with VAT followed Michaelis-Menten kinetics ^1^, with little cooperativity for ATP hydrolysis. This low cooperativity is inconsistent with simultaneous hydrolysis at all twelve nucleotide binding sites but is consistent with the proposed hand-over-hand mechanism, where only one NBD needs to bind and hydrolyze nucleotide at a time. The sequential model for ATP hydrolysis is also consistent with the asymmetry of the complex and nucleotide binding sites observed here, and the homology between VAT and other AAA+ enzymes known to hydrolyze ATP in a sequential fashion^15,17^.

**Figure 4.**
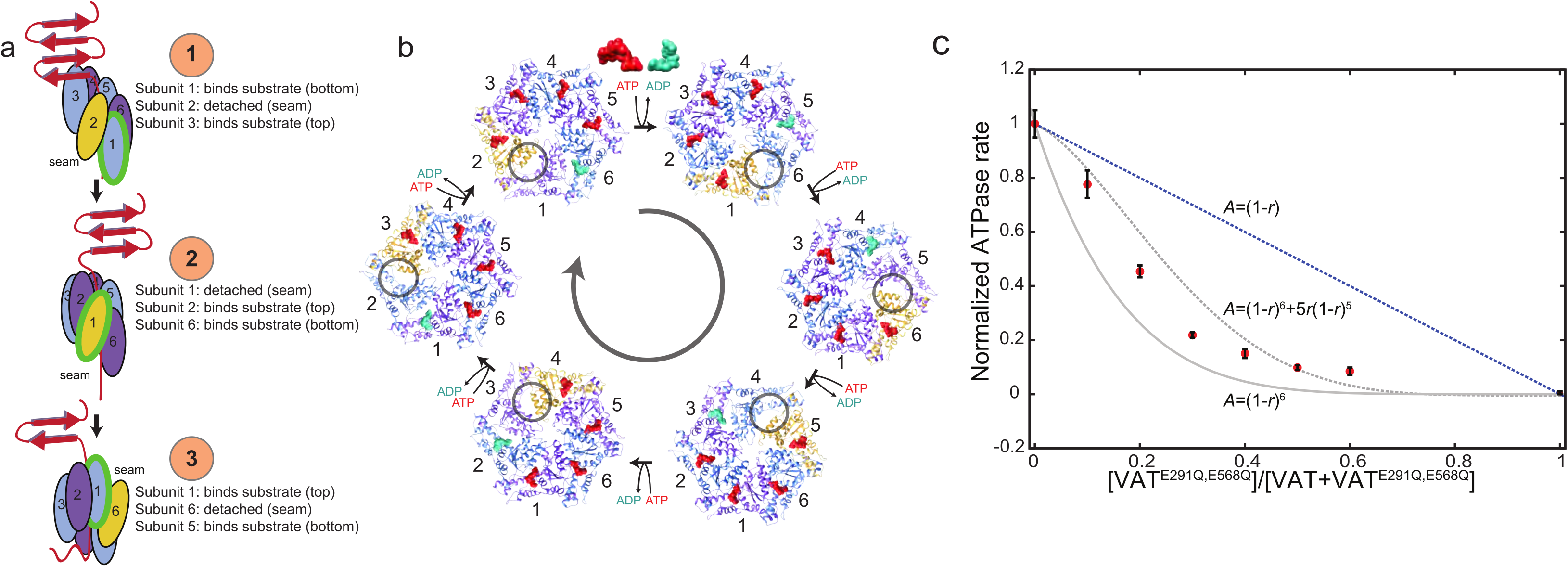
Model for substrate translocation by VAT. **a,** The substrate-bound structure of ΔN-VAT suggests that the enzyme is processive, with a single subunit (green outline) that binds substrate at the lowest position (step **1**) releasing as it becomes the seam subunit (yellow) (step **2)** before re-engaging further along the substrate (step **3)**. Subunits are numbered 1-6. **b,** The different conformations of VAT cycle around the hexamer, with ATP binding and ADP release with each transition. The seam subunit is indicated in yellow, with the adjacent subunits in the clockwise and counter-clockwise positions (viewed from NBD1 to NBD2) corresponding to protomers that are closest to and furthest from the folded substrate, respectively. **c,** The ATPase activity (*A*) of VAT with different ratios (*r*) of catalytically inactivated VAT^E291Q,E568Q^ mutant incorporated randomly into the enzyme hexamers. The activity decreases with increasing *r* faster than the *A* = (1-*r*)^6^+5*r*(1-*r*)^5^ curve expected if two inactive mutants per complex were required to stop VAT activity, instead approximating the *A* = (1-*r*)^6^ curve that is consistent with the proposed ‘hand-over-hand’ mechanism.

The cryo-EM structure of substrate-bound ∆N-VAT shows that both NBD rings of the complex adopt similar conformations (Fig. 3c and d). This similarity suggests that the mechanism of substrate unfolding proposed here may well operate in single-ring AAA+ unfoldases as well, including Rpt1-6 of the 19S proteasome, and the hexameric PAN and ClpX unfoldases. Thus, a picture emerges whereby AAA+ unfoldases adopt asymmetric substrate-bound conformations, creating narrow channels that sequester unfolded stretches of target proteins. Notably, even in molecular machines comprised of homo-oligomers, asymmetry can play an important functional role, in the case of VAT by priming the enzyme for processive translocation of its substrate.

## Statement of contributions

ZAR performed protein purification, electron microscopy, image analysis, atomic model building, and enzyme assays. RH created the ΔN-VAT construct and performed protein degradation assays. RA created the E291Q, E568Q VAT mutant and assisted with the purification of this protein. LEK and JLR supervised the research. ZAR and JLR wrote the manuscript with input from the other authors.

## Data deposition

Electron microscopy density maps are deposited in the Electron Microscopy Databank under accession numbers EMD-XXXX and EMD-XXXX. Atomic models are deposited in the Protein Databank under accession number XXXX.

## Acknowledgements

ZAR was supported by a scholarship from the Natural Sciences and Engineering Research Council of Canada. LEK and JLR were supported by the Canada Research Chairs program. This research was funded by operating grants MOP-133408 (LEK) and MOP-81294 (JLR) from the Canadian Institutes of Health Research.

---

**Video 1. Conformational changes in VAT upon substrate binding.** Interpolation between the stacked-ring and substrate-bound states of VAT, shown as ribbon diagrams (left) and Cα positions for the pore loop residues (right). Substrate is shown in red. Scale bars 25 Å. Please view video as loop.

**Video 2. Conformational changes in VAT monomers upon substrate binding.** Interpolation between the stacked-ring and substrate-bound states of VAT, shown as a ribbon diagram for two VAT monomers. Substrate is shown in red. Pore loop residues are indicated in green. Scale bar, 25 Å. Please view video as loop.

**Video 3. The hand-over-hand model for substrate translocation.** Interpolation between states where each subunit sequentially goes through the different possible positions in the helix. Each promoter’s pore loop binds a section of substrate at the top of the helix and escorts that section toward the bottom of the image through five steps. On the sixth step the protomer releases the substrate and becomes the seam protomer, moving up and away from the central pore before reengaging at the top of the substrate. An arbitrary subunit is marked in green to help maintain orientation. The structures are shown as ribbon diagrams (left) and Cα positions for the pore loop residues (right). Scale bars, 25 Å. Please view video as loop.

## Materials and Methods

### Protein expression and purification

An expression vector for the *T. acidophilum* VAT gene with the N-terminal 181 residues deleted (ΔN-VAT) ^5^ was prepared using a pProEx expression plasmid (Invitrogen). The construct included an N-terminal His6-tag and a tobacco etch virus (TEV) cleavage site between the His6-tag and the protein sequence. Expression and purification of ΔN-VAT were performed following the protocol described previously for full-length VAT^5^. Walker B (catalytically inactivating) mutations ^16^ in VAT were made in a pET 28b vector that encoded codon-optimized VAT with an N-terminal His6-tag and a tobacco etch virus (TEV) cleavage site between the His6-tag and the ΔN-VAT sequence. The mutant protein was purified in the same way as wild type VAT and ΔN-VAT ^5^. To produce mixed complexes of wild type and catalytically inactivated VAT, different ratios of wild type and mutant protein were mixed and diluted to 15 µM (monomer) in unfolding buffer (50 mM HEPES pH 7.5, 200 mM NaCl, 6 M guanidine hydrochloride), and then refolded into hexamers by fast dilution at a ratio 1:20 (v:v) into a refolding buffer (50 mM HEPES pH 7.5, 200 mM NaCl, 0.5 M arginine). After concentrating with a centrifuged concentrating device (Amicon), refolded VAT mixtures were subject to size exclusion chromatography and the hexamer fractions were collected. Little or no aggregate or monomeric VAT was detected after refolding. The ADP- and ATPγS-bound forms of ΔN-VAT were prepared by incubation overnight at 25 °C with apyrase (New England Biolabs; 0.1 U/mg of ΔN-VAT). Apyrase was then removed by size exclusion chromatography and nucleotide was added to 5 mM.

### Electron cryomicroscopy

Purified ΔN-VAT at ~20 mg/mL in buffer (50 mM HEPES pH 7.5, 100 mM NaCl) was incubated with 5 mM ATPγS (90 % pure, Sigma) for 5 min at room temperature before preparing cryo-EM grids. Immediately before grid freezing 0.05 % (w/v) IGEPAL CA-630 (Sigma-Aldrich) was added to the protein solution to increase the number of particles adopting side views on the grid. Sample (2.5 µL) of was applied to nanofabricated holey gold grids ^18–20^, with a hole size of ~1 µm and blotted using a modified FEI Vitribot for 5 s before plunge freezing in a liquid ethane/propane mixture (ratio of ~3:2) held at liquid nitrogen temperature ^21^. Micrographs were acquired as movies with an FEI Tecnai F20 electron microscope operating at 200 kV and equipped with a Gatan K2 Summit direct detector device camera. Movies, consisting of 30 frames at 2 frames/s, were collected with defocuses ranging from 1.7 to 2.9 µm. Movies were collected in counting mode with an exposure of 5 electrons/pixel/s, and a total exposure of 35 electrons/Å^2^.

### Electron microscopy image processing

Whole frame alignment of movies was performed with *alignframes_lmbfgs* ^22^ and the resulting averages of frames were used for contrast transfer function (CTF) determination with *CTFFIND4* ^23^ and automated particle image selection with *RELION* ^24^. Particle co-ordinates were used to extract particle images in 256 × 256 pixel boxes from the unaligned movies, while performing individual particle movement correction and exposure weighting with *alignparts_lmbfgs* ^22^. Magnification and CTF parameters were corrected for a previously measured 2 % magnification anisotropy ^25^ resulting in a calibrated pixel size of 1.45 Å. A total of 171,381 candidate particle images were subjected to two rounds of 2D classification, ignoring the CTF until the first peak, in *RELION* ^26^. Visual inspection and selection of classes yielded 93,469 particle images, which were then transferred to the program *cryoSPARC* ^9^ for *ab initio* 3D classification and refinement. Multiple 3D reference-free classifications were performed with between two and six classes using the stochastic gradient descent algorithm, before selecting two classes that yielded the highest quality maps. For refinement, a class distribution threshold of 0.9 was used to define individual particle images as belonging to a class. Refinement of the six-fold symmetric class of 75,205 particle images using C6 symmetry yielded a reconstruction at a resolution of 3.9 Å as measured by Fourier shell correlation (FSC) following a gold-standard refinement. A gold-standard refinement of the substrate-engaged class of 13,238 particle images with no symmetry applied yielded a reconstruction at a resolution of 4.8 Å by FSC.

### Map analysis and model building

Most of the density for loops, β-sheets and α-helices in the 3.9 Å resolution map were of sufficient quality to allow model building. A homology model of VAT that had previously been fit flexibly into a lower-resolution map ^5^ was rigidly fit into the map as individual domains (NBD1 and NBD2) with *UCSF Chimera* ^27^. Final models were built with successive rounds of manual model building in *Coot* ^28^ and real space refinement in *Phenix* ^29^, and gave an *EMringer* score of 1.17, which is slightly better than the typical score of 1.0 for a map at this resolution. For the substrate-bound complex, the resolution was insufficient to determine side chain orientations, and a refined structure from the symmetric class was fit into the density by molecular dynamics flexible fitting (*MDFF*^30^). Density for each monomer was segmented with *UCSF Chimera* ^27,31^. Only backbone atoms were subject to the fitting force during simulation, and optimization of geometry and rotamers was performed with *Phenix* ^29^.

### Degradation Assay

The auto-degradation of ΔN-VAT was measured following the protocol of Sauer and coworkers^4^. Hexameric ΔN-VAT (0.3 µM) was mixed with an ATP-regenerating system (20 U·mL-1 pyruvate kinase; 15 mM phosphoenolpyruvate) in the presence or absence of 10 mM ATP, with or without 0.9 µM *T. acidophilum* 20S proteasome. The reactions were carried out at 45 °C with a total reaction volume of 50 µL (50 mM HEPES pH 7.5, 200 mM NaCl, 120 mM MgCl2). Samples were withdrawn from the reaction at different times, quenched by heating in SDS-loading buffer, and subjected to SDS-PAGE. Intensities of the ΔN-VAT bands were analyzed with *Image Lab 4.0.1*.

### ATPase Assays

Steady state ATP hydrolysis was measured with an enzyme-coupled assay described previously ^32^. The reaction was carried out at a VAT concentration of 2.5 µM (monomer), and included an ATP regeneration system that contained 2.5 mM phosphoenolpyruvate, 0.2 mM NADH, 50 µg/mL pyruvate kinase, 50 µg/mL lactate dehydrogenase, 3 mM ATP, 200 mM NaCl, 120 mM MgCl2, and 50 mM HEPES (pH 7.5). VAT complexes were prepared as mixtures of wild type and inactivated subunits (Walker B mutations) where Glu291 (NBD1) and Glu568 (NBD2) were substituted with Gln, followed by the unfolding/refolding protocol described above. Hydrolysis of ATP was monitored in a UV-visible light spectrometer at 340 nm, 25 °C, in the presence or absence of 30 μM GFP substrate with an 11 residue ssRA tag ^4^. All measurements were done in triplicate. Similar results were obtained in the absence of substrate.

The expected activity of VAT complexes prepared by mixing different ratios of wild type and mutant protomers can be calculated as described previously ^16^. We define 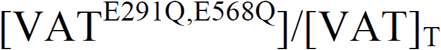, which can be calculated from the concentration of mutant VAT protomers that are mixed with wild type, where VAT^E291Q,E568Q^ is the VAT monomer bearing the E291Q/E568Q mutations and 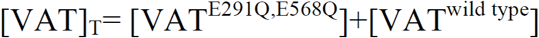. We assume that the unfolding/refolding process leads to equal probabilities of insertion of either a mutant or wild type protomer into a VAT hexamer, so that *r* is the probability of any protomer in the hexamer being a mutant. Under this assumption the fraction of VAT molecules with *k* mutant protomers is given by

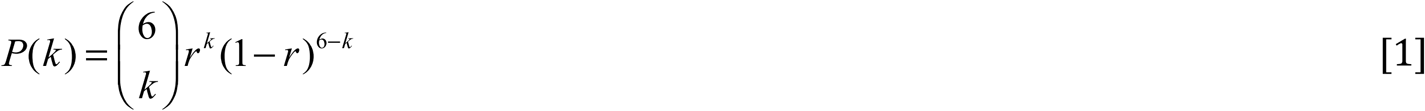

where 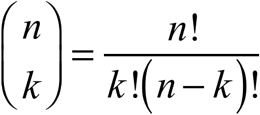, with the concentration of functional ATP hydrolysis sites in the protomer given by 2×(6-*k*) (two nucleotide sites per VAT). Assuming further that ATPase activity in the functional sites is not affected by the incorporation of inactive protomers until *m* are added, in which case all hydrolysis stops, one can write the fractional activity of a solution of VAT complexes as a function of *r*, as

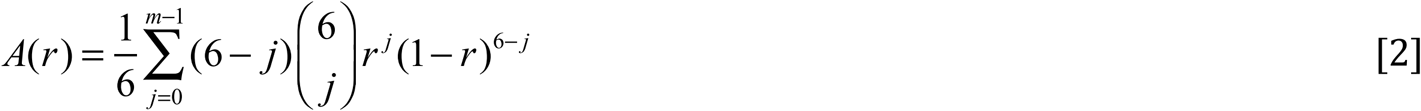

where *A*(*r*) is activity normalized relative to the expected hydrolysis rate for fully wild type VAT hexamers at an identical concentration. Our structural model predicts that a single defective protomer is enough to eliminate ATPase activity in the hexamer. In this case *m*=1 and *A*(*r*) = (1-*r*)^6^. In contrast, in the case where there is no coupling between active and inactive protomers,

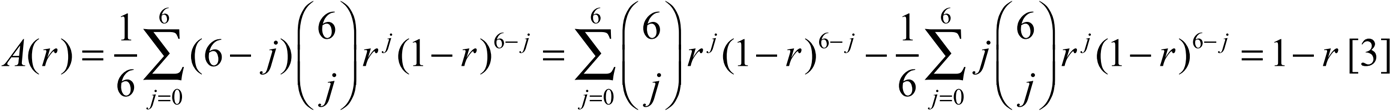

Figure 4c plots measured *A*(*r*) vs *r*, as obtained experimentally, showing that ATP hydrolysis rates decrease rapidly with *r*. Although the dependence of *A* on *r* is slightly less steep than expected for a model where a single defective protomer completely eliminates ATP hydrolysis (see text) the experimental data points decrease more steeply than predicted for *m*=2 (two defective protomers are required to eliminate hydrolysis).

**Figure 1 Supplement 1.**
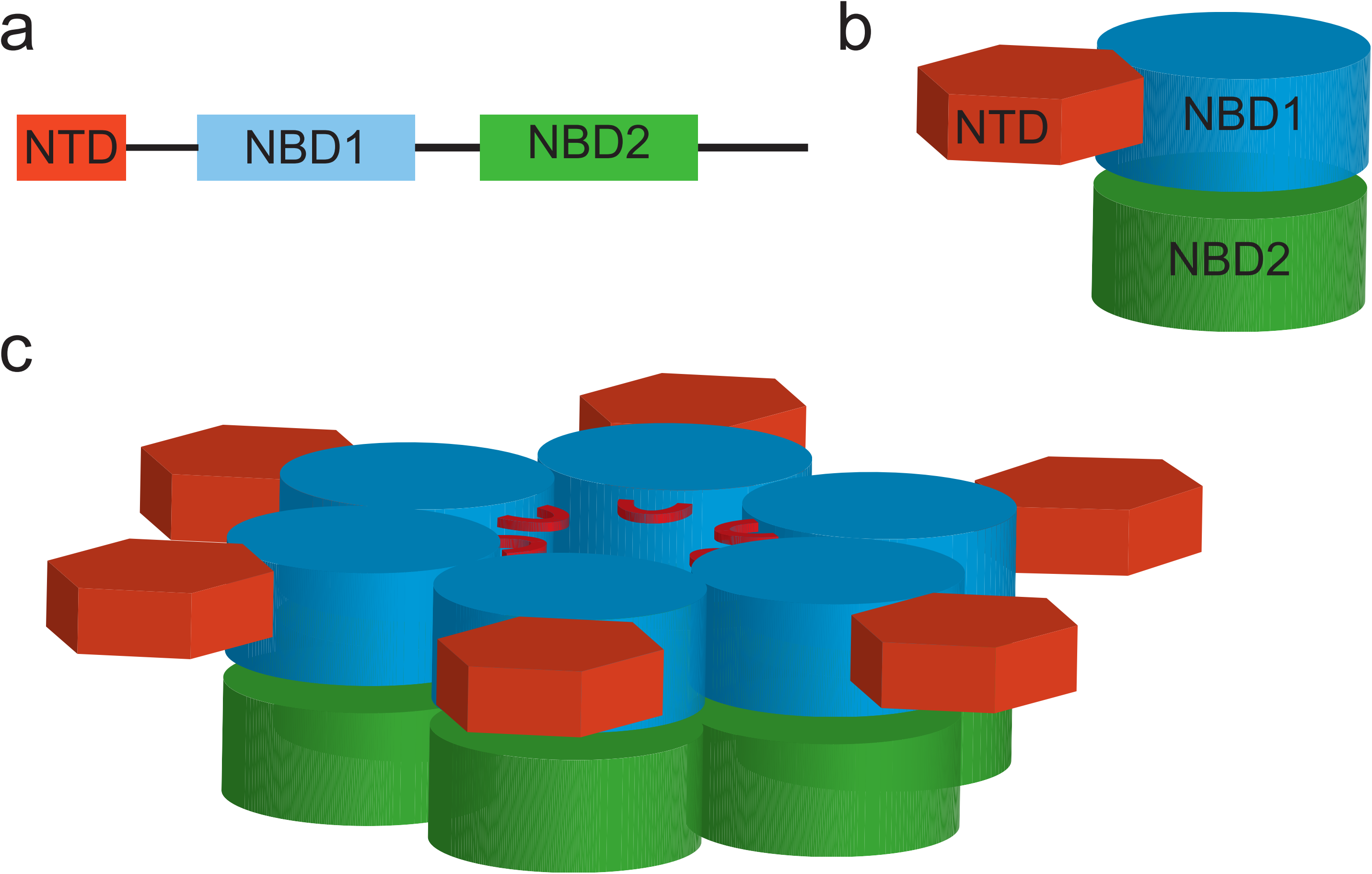
Domain arrangement and oligomeric structure of VAT. Cartoon representation of the VAT primary structure **(a)**, monomer structure **(b)**, and hexamer structure **(c)**. In **c** the stacked ring structure is shown with the AAA+ nucleotide binding domains colored in blue and green for NBD1 and NBD2, respectively, with the N-terminal domain (NTD: orange) co-planar with the NBD1 rings.

**Figure 2 Supplement 1.**
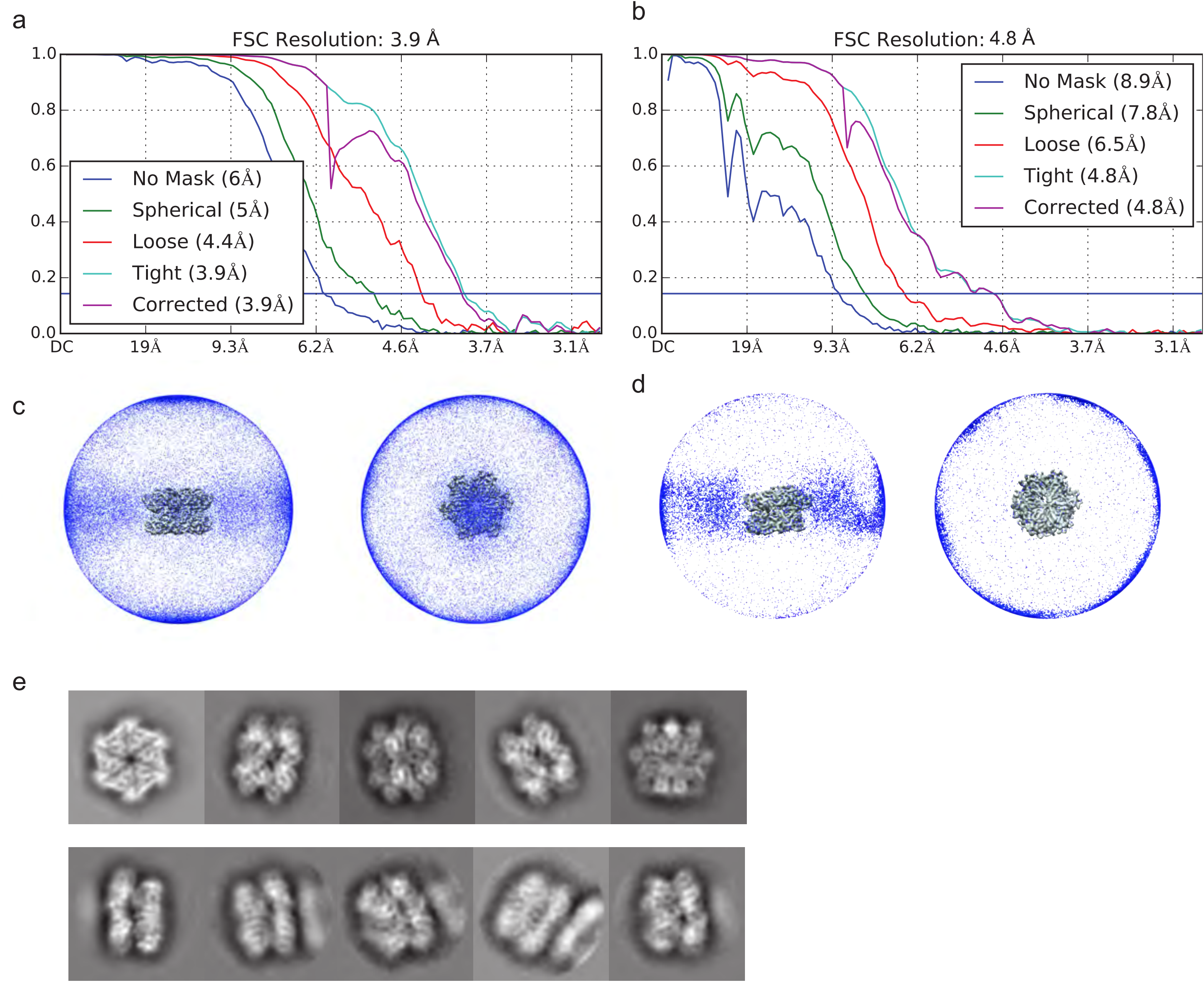
Cryo-EM map calculation and validation. **a,** Fourier shell correlation (FSC) curve for stacked-ring conformation of ΔN-VAT after a gold-standard refinement (resolutions reported at FSC=0.143). **b,** FSC curve for substrate-engaged conformation of ΔN-VAT after a gold-standard refinement (resolutions reported at FSC=0.143). **c,** Image orientation distribution for stacked-ring conformation. **d,** Image orientation distribution for substrate-engaged conformation. **e,** Representative 2D class average images for ΔN-VAT.

**Figure 2 Supplement 2.**
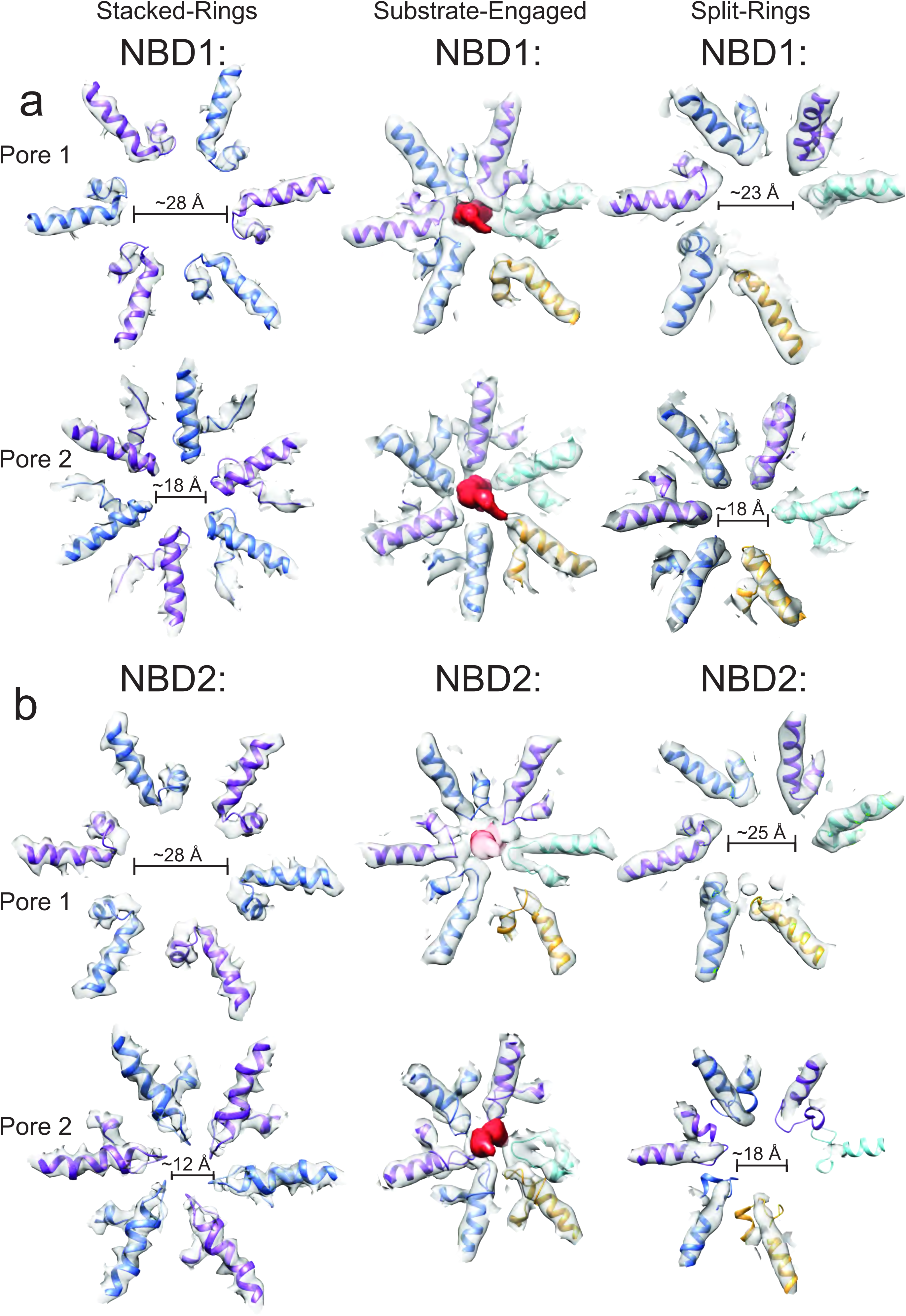
Pore loops of VAT. Compared to the stacked-ring conformation (left) and the split-ring conformation determined previously^5^ (right), the channel through the lumen of the substrate-engaged state of VAT (middle) formed by pore loop residues in both NBD1 **(a)** and NBD2 **(b)** is much more constrained. Note that the pore loops of the seam protomer do not participate in forming the channel (gold). Scale bar, 25 Å.

**Figure 3 Supplement 1.**
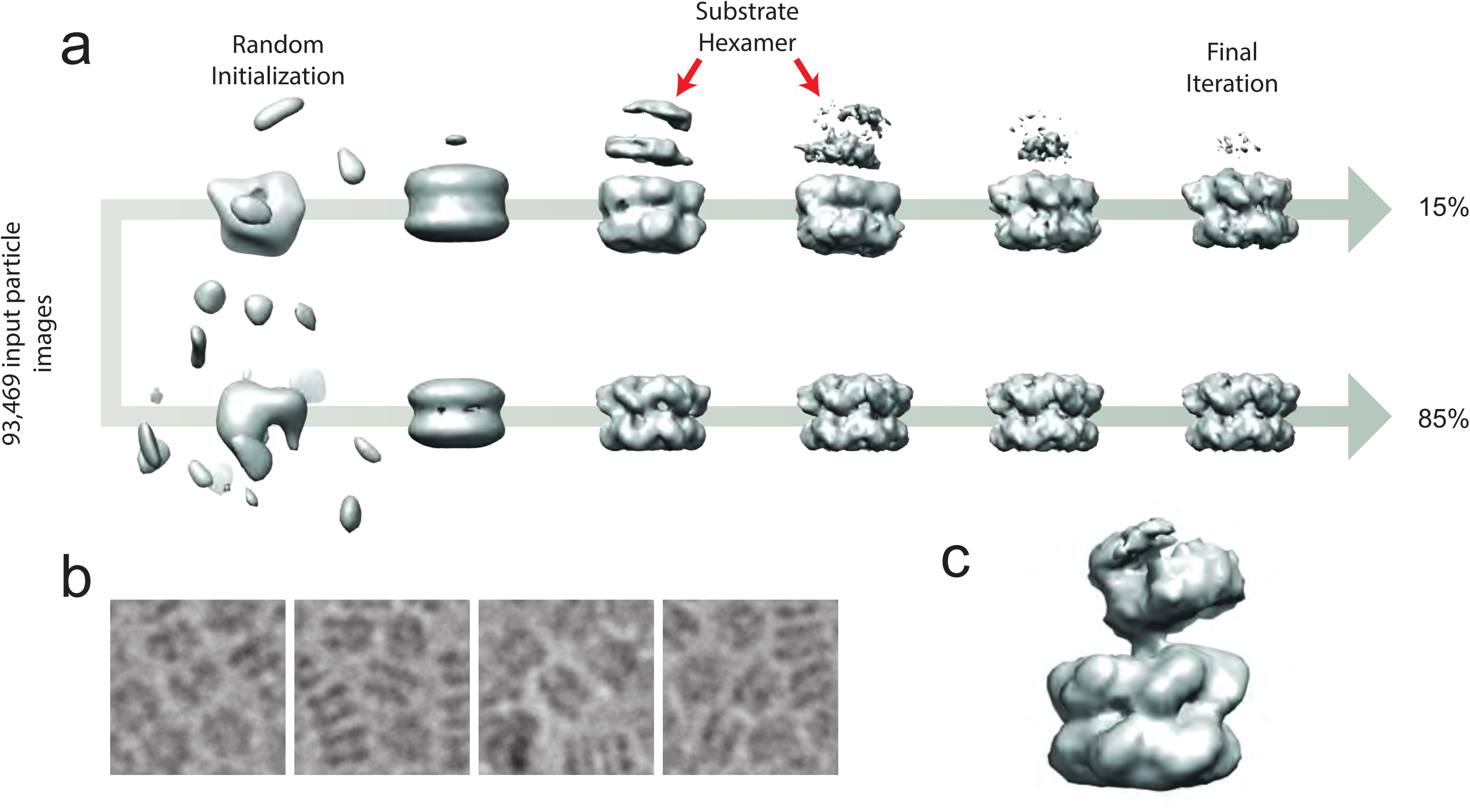
*Ab initio* classification to separate the six-fold symmetric and substrate-engaged states of VAT. **a,** 3D maps from the two classes throughout the *ab initio* classification scheme. At iteration ~500, the low-resolution shape of the substrate is apparent in the substrate-bound class (red arrows) but as the algorithm progresses, the density of this region decreases due to image alignment to the active VAT complex. The final distribution of single particle images was ~15 % substrate-engaged and ~85 % stacked-ring. **b,** Individual particle images showing crowding of the complexes. **c,** A low-pass filtered map of the substrate-engaged state of VAT shows the partially-folded substrate clearly.

**Figure 3 Supplement 2.**
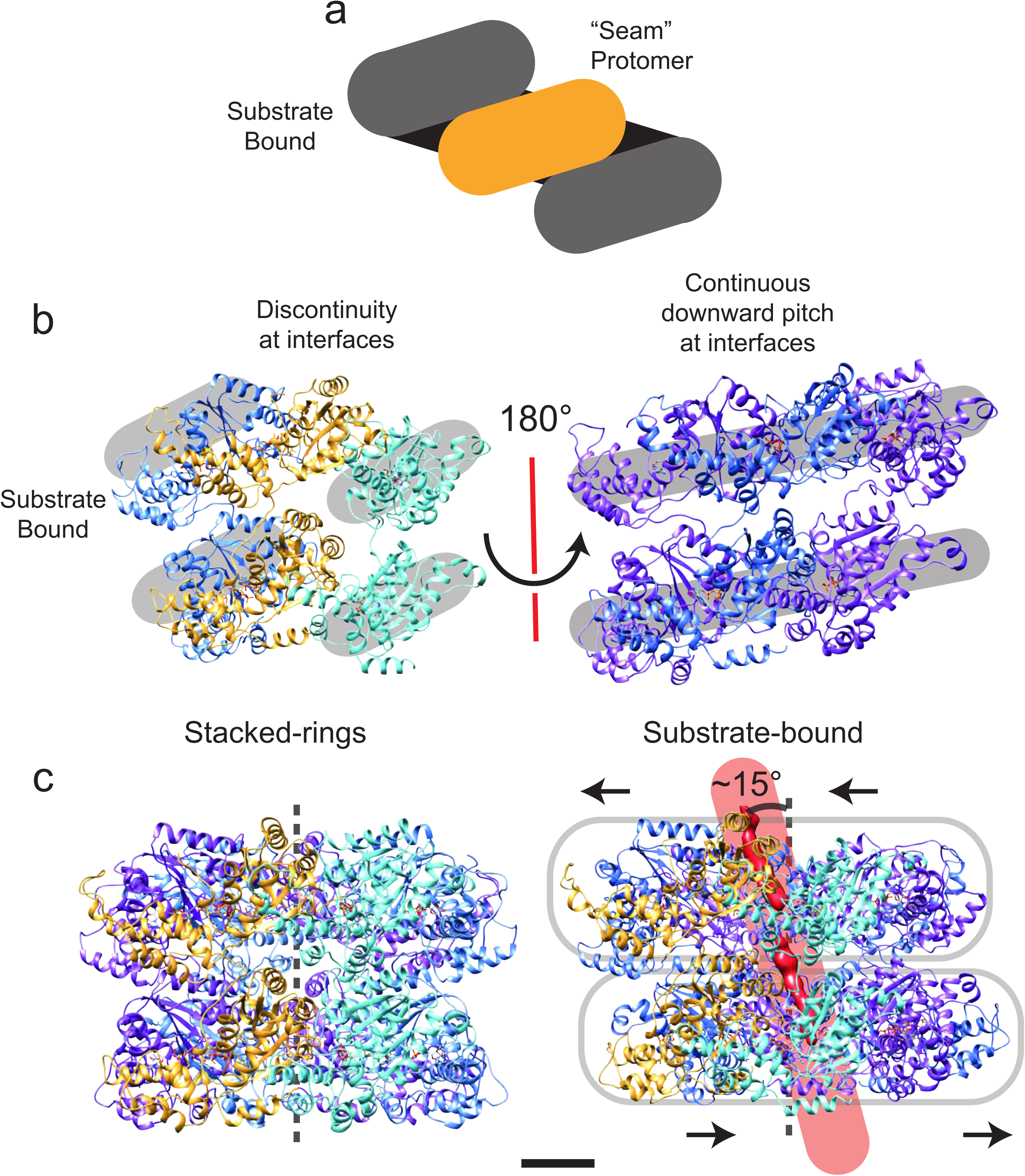
Symmetry breaking and seam formation in the AAA+ rings of substrate-engaged VAT. **a,** Cartoon showing the helical arrangement of AAA+ subunits bridged by a single subunit at the seam (gold). **b,** Views of the different types of protomer-protomer interfaces at the seam (left) and opposite the seam (right). The broken helix structure creates discontinuities between the lowest protomer (cyan) and the seam protomer (gold), as well as the highest protomer (blue) and the seam protomer (gold). **c,** Comparison of the central pore axis for stacked-ring (left) and substrate-engaged (right) states. Due to translation of the NBD1 and NBD2 rings relative to each other and tilting of the pore, the central pore axis passes through the complex at an angle of ~15°. Scale bars, 25 Å.

